# On doubting image quality assessment metrics for microscopy virtual staining

**DOI:** 10.64898/2026.08.31.748410

**Authors:** Wei-shan Li, Gregory P. Way

**Affiliations:** Department of Biomedical Informatics, University of Colorado Anschutz, Aurora, CO

## Abstract

Pairing label-free microscopy with virtual staining could reduce the cost and experimental burden of fluorescence microscopy, but its impact is conditional on generalizable inference. Most virtual staining studies assess performance using image quality assessment (IQA) metrics developed for natural images, yet how well these metrics translate to microscopy remains unknown. Here, we examined the behavior of seven commonly-used full-reference training objectives and metrics, MAE, PSNR, SSIM, foreground PSNR and SSIM, LPIPS, and DISTS, under controlled image degradation and realistic out-of-distribution virtual staining. We applied graded intensity, textural, and morphological transformations to Cell Painting images spanning 18 cell lines, seeding densities, and fluorescence channels. Channel, cell line identity and seeding density explained substantial metric variation after controlling for degradation magnitude. DISTS and foreground metrics showed more favorable balance between degradation sensitivity and biological invariance, although no metric reported performance independent of biological context. Incrementally degrading images and evaluating concomitant metric degradation further revealed that most metrics used only a small fraction of their nominal numerical ranges and frequently plateaued while image degradation visibly continued. We next trained three popular virtual staining model architectures (UNet, WGAN-GP, UNeXt) on five U2-OS seeding densities separately, and computed metrics on model predictions across 17 unseen cell lines. We observed that architecture and training U2-OS seeding density together explain less than 2% of metric variation. Visual inspection suggested comparable scores across cell lines correspond to qualitatively distinct errors, such as differences in cell morphology and marker intensity. These findings show that conventional IQA metrics do not effectively translate to virtual staining applications. Selection or optimization of virtual staining models against real application such as in label-free high content drug screening should instead be approached in an application-oriented fashion.

## Introduction

Deep learning-based virtual staining offers a potential alternative to fluorescence-based imaging assays by inferring labeled subcellular structures from label-free modalities such as brightfield or phase-contrast microscopy (1–5). If virtual staining models can faithfully learn generalizable mappings from label-free and non-destructive modalities to fluorescence channels, they can be applied virtually cost-free as substitutes to labor-intensive and destructive sample-preparation procedures (6,7). This opportunity is particularly relevant to Cell Painting and related multiplex fluorescence assays, which characterize cell state by marking multiple organelles and subcellular compartments in an unbiased fashion (8,9). The resulting morphology profiles can support analyses of drug mechanisms of action (MOA), phenotypic similarity, compound prioritization, and many other applications, but the staining and assay-optimization procedures required to generate them remain important constraints on throughput, cost, and interpretability (8–12). Most cell microscopy virtual staining research has focused on adapting increasingly-capable and innovative image-to-image translation architectures to label-free prediction of fluorescence stains, such as the vanilla UNet, generative adversarial network (GAN), and recent adaptation of the ConvNeXt vision backbone, alongside training strategies such as masked autoencoder pretraining to improve learning (2–4,13,14). The more advanced architectures were originally developed in natural-image computer vision fields, which model pictures or videos of the natural world like animals, cars and landscape. Recent studies have established that these architectures can accurately predict fluorescence microscopy images under stable data distributions.

Despite these advances, an important question connecting modelling success to real application in large-scale image-based profiling remains insufficiently addressed: how reliable are label-free virtual staining models on out-of-distribution (OOD) cell microscopy data? Published evaluations use held-out images acquired within the same experimental system and imaging protocol as the training data with limited variation in biological context. While this design quantifies interpolation across data batches within single institutions, it informs little extrapolation performance in profiling applications such as a drug screen where models must predict not only unseen cell types, but also unseen perturbations and cell seeding densities acquired across different imaging platforms (15). Models trained on narrow biological contexts may perform well under conventional heldout validation yet fail under a complex OOD cell state (16,17). A related challenge concerns how practitioners measure virtual staining performance. Most studies rely on image quality assessment (IQA) metrics such as peak signal-to-noise ratio (PSNR), structural similarity (SSIM), learned perceptual image patch similarity (LPIPS), and deep image structure and texture similarity (DISTS) (18,19). Like model architectures, most image metrics were developed for natural-image applications and guarantee neither content-invariant comparisons across heterogeneous images nor intuitive interpretations (20). Consequently, metrics values may reflect not only reconstruction fidelity but also on variation in biological conditions like cell morphology, cell density, and cell type. Realizing the promise of virtual staining therefore requires establishing whether the metrics used to measure performance remain interpretable across the relevant sources of biological variation.

In this study, we answer two related questions. First, how do widely used IQA metrics respond to controlled image degradations across biologically heterogeneous image content? Second, through the lens of natural image IQA metrics, how do contemporary virtual staining models compare across biologically-relevant OOD variation? To answer these questions, we analyze a diverse Cell Painting dataset of 18 pediatric cancer cell lines assayed across five seeding densities and two experimental plates. We create a synthetic dataset by applying six types of controlled image degradations (e.g., erosion, Gaussian noise, etc.) in six increasing steps of severity gradations to these microscopy images. Using regression-based variance decomposition, we characterize the sensitivity, robustness, and susceptibility to confounding by biological variation commonly used IQA metrics in virtual staining. We next investigate model generalization by training three representative virtual staining architectures: UNet, UNet augmented with a WGAN-GP discriminator, and a simplified 2D UNeXt (2,4,13). We intentionally train these model architectures under a highly-restricted domain consisting of 450 full FOVs of untreated U2-OS cells, plated under specific seeding densities, from the first experimental plate. We evaluate these models across 17 additional unseen cell lines across two plates. This design simulates a scenario applying virtual staining beyond the narrow biological conditions available during training. We found that IQA scores were strongly influenced by biological context, frequently occupied only a restricted fraction of their nominal numerical ranges, and could lose discriminatory power while visible image degradation continued. We also observed that biological variables continued to explain more metric variation than model architecture or U2-OS seeding density used for training, and comparable scores could correspond to qualitatively different reconstruction failures. Taken together, our results demonstrate that commonly used IQA metrics are useful for detecting broad performance shifts but do not provide a context-invariant scale of biological reconstruction fidelity, motivating evaluation frameworks to take extra caution in interpreting metrics across OOD data, and better yet fully adopt application-oriented biological validation.

## Results

### We analyze a diverse Cell Painting dataset of pediatric cancer cell lines

Computing full reference IQA metrics requires paired reference images and altered counterparts whose differences can be evaluated under known biological contexts. All of our metric analyses operates around the same reference image set, which is a Cell Painting dataset representing 18 different pediatric cancer cell lines each plated under five discrete levels of seeding densities (1k, 2k, 4k, 8k, and 12k per well) (**Figure 1A**). Because the combination of 18 pediatric cancer cell lines and five seeding densities yielded 90 biological conditions, we name this dataset PedCP90. The PedCP90 dataset is collected from two 384-well plates, in a single image acquisition batch. Following image quality control, 10,248 Cell Painting FOVs remain, totalling to 176GB. All FOVs contained the five canonical Cell Painting fluorescence channels paired with a single-plane brightfield channel. Besides U2-OS which appears on both plates, all other cell lines appear on only one of the two plates **(Supplementary Figure 1**). The dataset is collected to include real biological variation in cell line, density, morphology, and plating differences while limiting technical variation from image acquisition. From PedCP90, we performed nuclei segmentation and obtained 18,818 nuclei centered crops from 10,248 full FOVs. From the cropped PedCP90, we generated two “altered” image sets for complementary characterization of metric behavior, PedCP90Sim and PedCP90VS.

**Figure 1:**
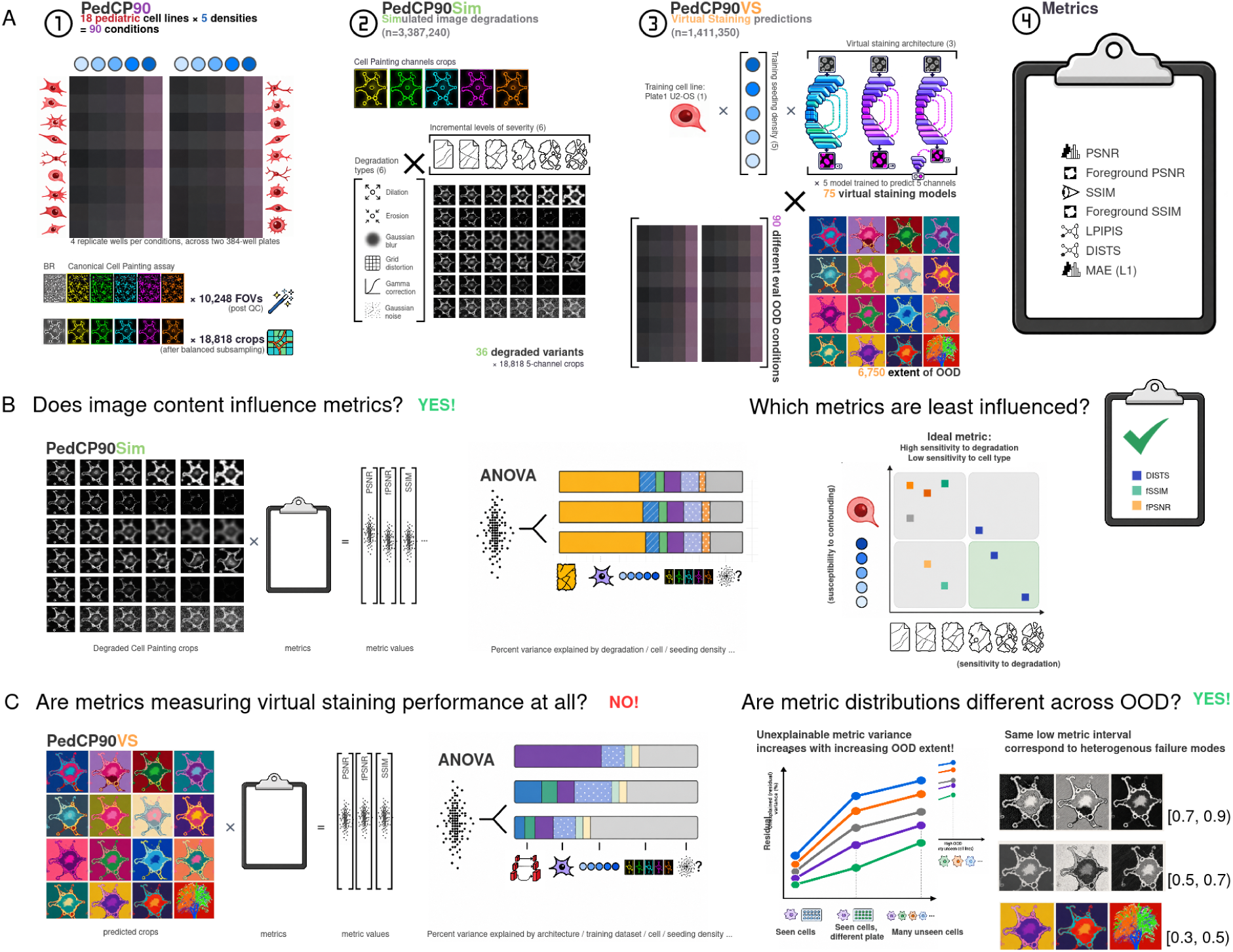
Schematic outlining our virtual staining metric focused analysis. (**A**) PedCP90 is a Cell Painting dataset containing 18 pediatric cell lines plated at five seeding densities. We derive two additional datasets, PedCP90Sim and PedCP90VS, from PedCP90. PedCP90Sim is a simulated image dataset created by adding six types of controlled degradations each at six graded severities to real Cell Painting images. PedCP90VS is a virtual staining OOD prediction dataset containing model predictions from three architectures each trained at five training seeding densities, on unseen cell lines. In this work, we apply image-quality assessment (IQA) metrics to PedCPsim and PedCP90VS using the reference PedCP90 as reference and analyze metric response to simulated signal and data-inherent variations in biology. (**B**) Analyzing PedCP90Sim. Metrics are computed per degradation type, and analyzed with ANOVA against data-inherent variations and simulation signals to quantify influence of these factors on metric values. Our results indicate that most metrics are heavily influenced by content of the image and the biology that drives such content difference. We report that DISTS and the foreground metric variants are relatively less susceptible to that influence. (**C**) Analyzing PedCP90VS. Metrics are computed on the OOD prediction images. ANOVA found that model architecture and training density had negligible impact on metrics and biology and residual variance still dominates metric variation. Further quantitative and qualitative analysis revealed increasing residual variance with increasing extent of OOD, and similarly distributed metric values can correspond to vastly different failure modes as evaluation context shifts away from train.

The PedCP90Sim assesses metric response under controlled, interpretable, and singular image error types while PedCP90VS assesses metric behavior against realistic virtual staining imperfections, where the severity and nature of error are both unknown. We describe each data variant briefly here, but see the **Methods** and for more details. For PedCP90Sim, we applied graded intensity, texture, morphology, distortion, and noise transformations to all real PedCP90 fluorescence images (**Figure 1A**). Because the transformation type and severity are known by construction, PedCP90Sim allows subsequent analysis of metric response to simulated reconstruction error type and extent in the midst of the 90 real biological variations (**Figure 1B**). For PedCP90VS, we used paired brightfield images from cropped PedCP90 images of the U2-OS cell line to train three representative virtual staining architectures per cell seeding density to predict the corresponding Cell Painting fluorescence channels (**Figure 1A**). This procedure resulted in 15 distinct models predicting each Cell Painting channel. We then applied virtual staining models to heldout U2-OS images and on biologically-distinct cell lines and seeding density conditions, producing 15 “altered” versions per single-reference fluorescence image in PedCP90. PedCP90VS complements PedCP90Sim by providing a limited yet more realistic testbed for measuring if metrics can discern architecture to architecture differences in the presence of the 90 biological variations (**Figure 1C**).

We applied seven IQA metrics commonly used in virtual staining to both PedCP90Sim and PedCP90VS. We selected four full-reference IQA metrics: peak signal-to-noise ratio (PSNR), structural similarity index measure (SSIM), learned perceptual image patch similarity (LPIPS), and deep image structure and texture similarity (DISTS) (**Supplementary Table 2**). Motivated by a recent, foreground-aware virtual staining approach called Spotlight reporting reduced artifacting and altered downstream morphology measurements by training virtual staining models on foreground prediction exclusively, we also created foreground-masked variants of PSNR and SSIM, limiting computation of these metrics to the Otsu foreground of the reference image (21). Lastly, we included mean absolute error (MAE), also known as L1, as it is the predominant objective function for training virtual staining models and is a perfectly unambitious pixel-wise error measure. We are primarily interested in whether the conventional metrics are influenced by image to image variation and biology, and whether foreground masking alters metric behavior in a useful way.

### IQA metrics are disproportionately sensitive to degradations across biological variations

An ideal metric for evaluating virtual staining performance should capture the error between model prediction and ground-truth reference, while ignoring inherent biological variation such as cell type, cell state, and cell density. We hypothesize that biological variations bias all metrics, but some metrics may be less susceptible to biological confounding. To test this hypothesis, we created the PedCP90Sim dataset by applying a suite of six image-degrading transformations at six incremental levels of severity to 18,818 image crops across all Cell Painting channels from PedCP90. The seven degradation transforms probe qualitatively distinct classes of image reconstruction failure modes. Briefly, we apply two morphological operators, dilation and erosion, to mimic morphology-altering errors such as over-expanded or under-recovered cellular structures. We apply Gaussian blur to represent loss of textural sharpness, which is a known common artifact of virtual staining (17,22); Grid distortion as a spatial deformation perturbation to warp object geometry locally while minimizing alteration of intensity distribution; Gamma correction (from brightening and dimming), to provide a complementary intensity-only perturbation monotonically changing only the pixel intensity distribution and range; and lastly, Gaussian noise to serve as a reference, positive-control perturbation that we expect all metrics to detect as it introduces additive noise independent from both morphology information and pixel intensity distribution. For each image-degrading transformation, we selected one primary parameter controlling for degradation severity and defined six incremental degrading levels, chosen to span a graded range from mild to severe degradation within each transformation type (**Figure 1A**; **Supplementary Table 3**). See **Methods** for more details.

We begin by applying ANOVA (type II) per metric and degradation type combination, as it simultaneously quantifies variance explained by cell line, seeding density, channel, and degradation severity in PedCP90Sim, providing holistic insights on what factors influence metrics the most (**Figure 1B**). We first report and interpret degradation specific trends regarding sensitivity to severity observable consistently across metrics. Gaussian noise is consistently the easiest to detect, explaining around 60%-90% of the variation in metrics (**Figure 2A**). Gaussian blur is the most difficult to measure, explaining only 10%-20% of metric variation. Interestingly, gamma correction, which alters pixel intensities to make darker pixels appear brighter or vice versa, received mixed sensitivity across metrics, explaining up to 60% of metric value variation in DISTS, LPIPS and SSIM, yet less than 20% in MAE, PSNR, and foreground PSNR. In terms of confounding effects, we observe, broadly, that the Cell Painting channels contribute the most to metric scores, accounting for 5%-30% of the variation (**Figure 2B**). The interaction term between channel and seeding density contributes 5%-10% variation, and cell line identity contributes between 0%-10% variation. Channel is the least concerning confounder, since the standard practice of stratifying virtual staining performance by channel eliminates its effect. Cell line and its interaction with seeding density pose a more serious limitation: the same degradation produces unequal metric changes across cell lines, which reflects biological differences in spatial structure and texture. Therefore, differences in selection of certain cell lines or drifts in confluence in the evaluation microscopy image can make the same virtual staining model appear artificially better or worse than it actually is.

**Figure 2:**
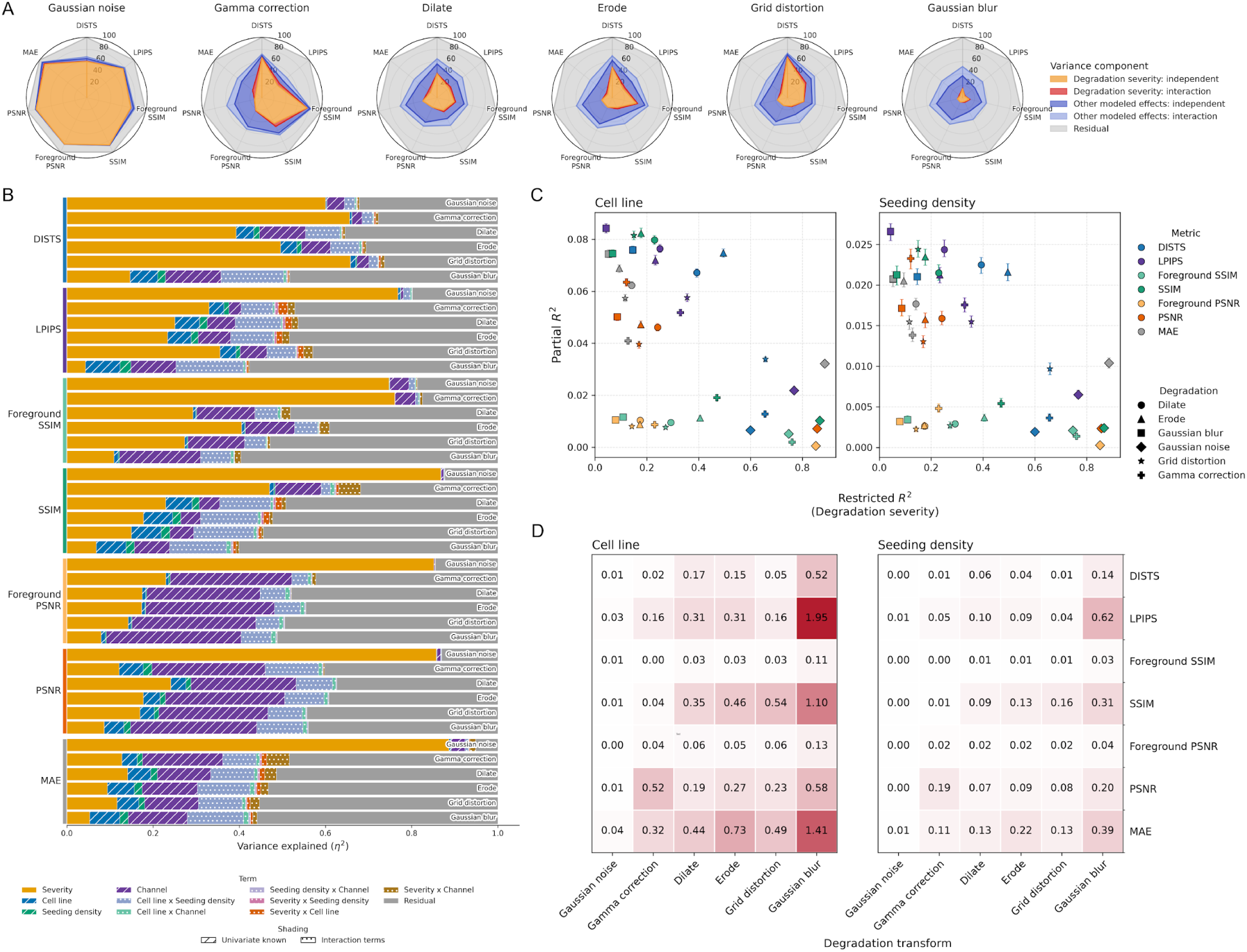
Metric susceptibility to confounding by biological variables in simulated degradation setting. (**A**) ANOVA analysis with explained variance terms aggregated across broad categories: univariate severity (yellow), severity interaction (red), univariate biological confounder (dark blue), biological confounder interaction (light blue), and residual (gray). Same analysis as B. (**B**) Full ANOVA decomposition of all variables, across metric and degradation. (**C**) Focused nested regression analysis results quantifying the relative contribution of severity against one of two dominant biological covariate at a time: Nested regression coefficient of determination analysis quantifying the respective contribution of biological context driven image content variation (bias, y axis) and known image transformation magnitude driven degradation (signal, x axis). Each point is a single nested regression analysis interrogating the response of one metric against one degradation type. Left column: confounder is cell line; Right column: confounder is seeding density; Upper row: colored by metric; Lower row: marked by degradation type. (**D**) Summary of analysis as metric burden (ratio of bias to sensitivity). Higher values (≥1) indicate metrics tend to measure content more so compared to capturing actual image degradation.

#### DISTS and foreground-masked metrics are influenced the least by biological confounding

Having established that metric sensitivity and biological confounding depend jointly on metric and degradation type, we next performed a focused bootstrapped nested regression analysis aimed at translating these observations into practical guidance for metric selection grounded by statistical confidence. For each regression formulation, the restricted model contained degradation severity alone, whereas the full model additionally included either seeding density or cell-line identity. The restricted R^2^ quantifies how strongly a metric tracks controlled degradation, while partial R^2^ quantifies the additional contribution of biological context after accounting for degradation severity. Within this framework, a desirable metric occupies the regime of high restricted R^2^ and low partial R^2^, changing substantially with reconstruction error while remaining comparatively stable across biological contexts (**Figure 2C**).

We observed metric performance that was largely consistent with the preceding analysis. DISTS maintained highest restricted R^2^ values across the spatial degradation classes, whereas other metrics were differentially sensitive across different degradations. We observed that the absolute contribution of seeding density was small, accounting for approximately 0-0.6% of metric variation in SSIM, PSNR, LPIPS, and MAE (**Figure 2C**). Restricting metric computation to reference-derived cellular foreground further improved metrics by decreasing influence by seeding density across degradation classes. For SSIM specifically, foreground restriction additionally resulted in increasing contribution by severity of erosion, dilation, and grid distortion, potentially making it a better metric more sensitive to decay in image quality. These paired comparisons point to foreground-to-background composition as an important mechanism through which cell density influences metrics: as confluence changes, the proportion of largely uninformative background pixels changes with it, altering metrics even when the underlying degradation severity is comparable.

We next examined the contribution of cell-line identity. Different cell lines can vary substantially in size, shape, texture, organelle organization, growth pattern, and fluorescence intensity distribution. Therefore, a metric that is sensitive to cell-line identity may assign different quality scores to equally degraded images simply because the underlying cellular content differs and gets differentially altered by the degradation. We observed that DISTS remained best in class. DISTS was influenced by grid distortion while limiting influence to cell line confounding. However, cell-line identity contributed to measurable variance, from 4 to 8.5%, across all metrics under erosion, dilation, and gaussian blur (**Figure 2C**). Foreground restriction again substantially reduced the contribution of cell-line identity for SSIM and PSNR, in several cases from approximately 4–8% to around 1%, placing the foreground variants among the least influenced by cell line confounding. Surprisingly, our results in the cell line point to the same confounding mechanism mediated by foreground-to-background ratio. The bulk of the cell line confounding seemed correctable by foreground masking, yet an unanimous ∼1% cell-line confounding effect seemed to persist in all foreground metric combinations, plausibly reflecting morphological and textural differences evident in the foreground alone.

To summarize the tradeoff between degradation sensitivity and biological confounding, we next computed a burden statistic as the ratio of partial R^2^ attributable to biological context to restricted R^2^ attributable to degradation detection (**Figure 2D**). Foreground SSIM and PSNR showed among the lowest burdens across transformation classes and both biological confounders, while DISTS combined low burden with comparatively strong sensitivity to several challenging degradations, particularly grid distortion and gamma correction. In contrast, several conventional metrics approached or exceeded a burden of 1.0 against non-trivial degradations such as Gaussian blur, erosion/dilation and grid distortion, to the point where biological context rivals or exceeds the intended degradation signal. Together, these results identify DISTS and foreground-masked metrics as the most consistently resistant to biological confounding, although their relative advantage remains failure-mode dependent.

#### IQA metrics values risk implying misleadingly high performance and unreliable model comparison

Our preceding analyses characterized how IQA metrics are influenced by controlled degradation and biological confounding. However, humans do not automatically comprehend IQA metrics as calculated fractions of explained variance. Instead, humans see and interpret the absolute metric values, for example, when comparing competing architectures. In other words, a metric may behave favorably in variance analyses yet be difficult or misleading to interpret on its native numerical range. We consequently asked a complementary question: as microscopy image fidelity progressively deteriorates, how intuitive is the absolute numerical response of an IQA metric?

For a metric to be interpretable, incremental degradation of the same image should produce correspondingly graded changes in score. A near perfect prediction should remain close to the metric optimum. Moreover, as degradation severity increases, deterioration in metric score should occupy enough of the available numerical range to distinguish different levels of prediction quality. We would also expect an eventual flattening of metrics with increasing degradation severity because the degradation removes increasingly marginal information. For example, further Gaussian blurring of an image that already lost all high frequency texture should not be expected to produce significant score change. To probe such fine-grained response trajectories from metrics while controlling for biological variation, we applied the transformation with very minimal severity setting at one-step increments for 500 iterations on a single sampled fluorescence image from PedCP90. For all metrics-degredation trajectories, we characterized the following properties: metric ceiling attainment, the empirical metric dynamic range, and trajectory plateaus (**Figure 3A**).

**Figure 3:**
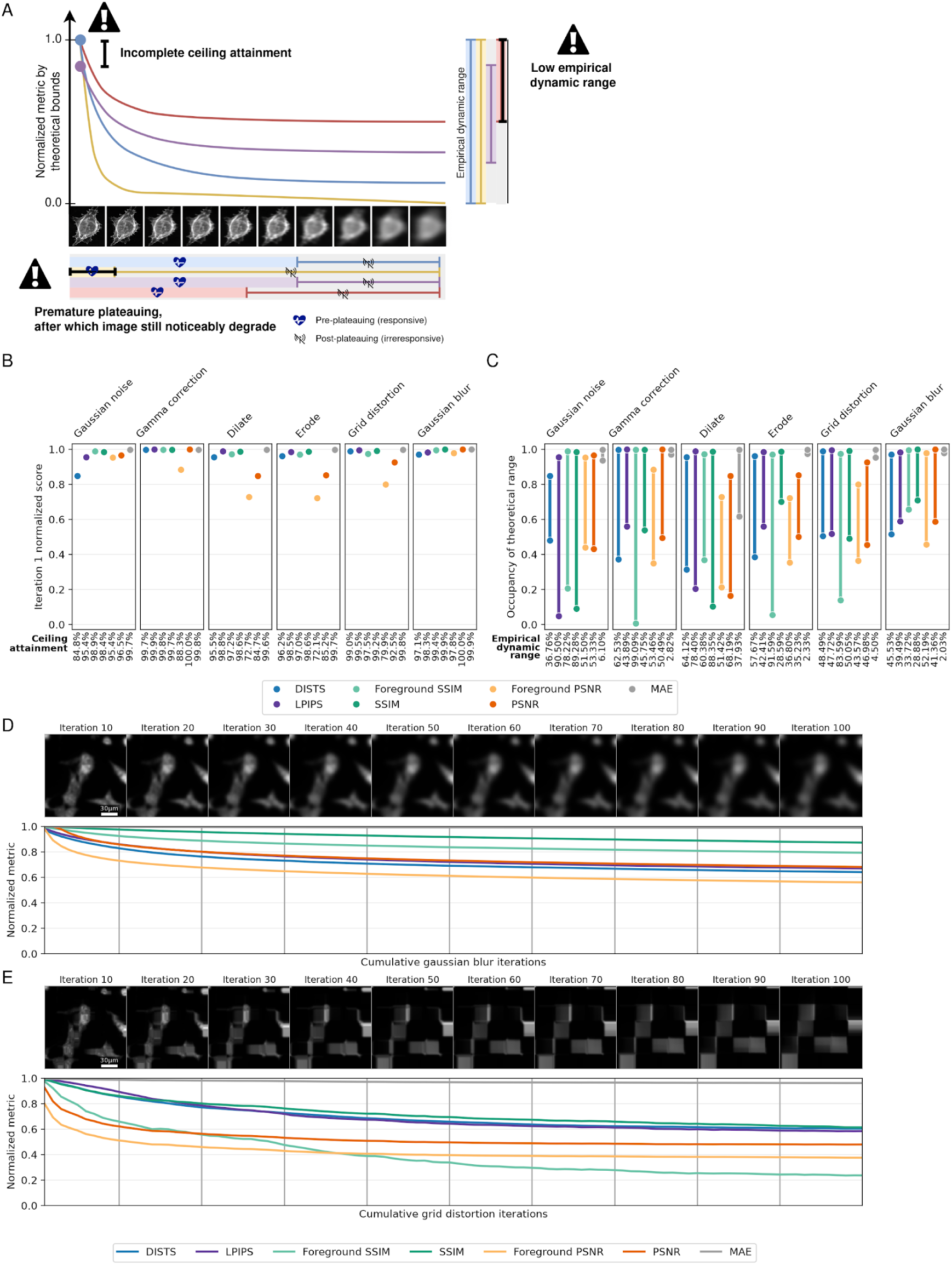
Metric absolute response against simulated degradation, controlling for biological context. **(A)** We define three properties characterizing metric absolute response to progressive image degradation: ceiling attainment, the fraction of the theoretical maximum reached under minimal degradation; empirical dynamic range, the fraction of the nominal metric range traversed across the degradation trajectory; and plateauing behavior, the point at which further degradation produces diminishing score changes. Low ceiling attainment can exaggerate minor errors, restricted dynamic range can make severe degradation appear deceptively favorable, and premature plateauing indicates loss of sensitivity to continued image deterioration. **(B)** Summary of ceiling attainment by metrics on iteration 1 **(C)** Summary of empirical dynamic range of all metrics against all degradation types. Metric responses and montages for the first 100 iterations against **(D)** Incremental gaussian blurring, **(E)** incremental grid distortion.

All metrics follow a clear exponential pattern where metric scores decay rapidly toward the high-quality end of the trajectories before transitioning into a plateauing behavior (**Supplementary Figure 2**). This decay pattern is ideal for virtual staining model evaluation as it concentrates discriminatory power (room for metric variation) to the range between perfect and near-perfection.

Ceiling attainment was generally favorable. Among metrics that have a clearly documented or mathematically derivable limit, namely DISTS, LPIPS, SSIM, and foreground SSIM, the first degradation step attained more than 97% of their theoretical ceiling value and stayed close to the ceiling attainment in terms of pure average pixel error (MAE) for most transformation classes (**Figure 3B**). Erosion and dilation produced larger initial changes relative to MAE because even their smallest configurable operation alters whole objects, while DISTS under Gaussian noise represented an additional outlier. Our findings indicate that the theoretical max of most natural image IQA metrics against most simulated degradation types are, in practice, attainable in cell microscopy images, suggesting that we can reliably compare the quality of virtual staining prediction should it be near the perfection range.

However, across most metric-degradation combinations, the degradation trajectory from minimal to severely corrupted traversed less than 50% of their nominal range (**Figure 3C**). In several cases, even the most severely degraded images retained normalized fidelity scores of approximately 0.5–0.6, suggesting that interpreting IQA scores with respect to their nominal range could misleadingly represent severely corrupted images as medium quality. The low observed restricted empirical dynamic range motivated a more consequential question to ask about metric behavior at and following plateauing. Does a metric plateau because further degradation no longer meaningfully changes the image, or because the metric has become arbitrarily insensitive while image quality continues to deteriorate? We refer to these two hypothesized plateauing behaviors as content-exhaustion plateaus and sensitivity-limited premature plateaus, respectively.

For pixel intensity adjusting degradations, metric plateauing seemed driven by content-exhaustion. For gamma correction, we observe a stable decay against the first 20 iterations where the image content remained but gets increasingly low contrast, followed by a plateau in most metrics starting iteration 40 where the repeated lowering of contrast effectively rendered the entire image blank and subsequent operations no longer create meaningful changes (**Supplementary Figure 3A**). Erosion behaved similarly and even more stably, with an initial metric descent exactly concordant with foreground removal and later flattening corresponding to the lack of noticeable subsequent erosion (**Supplementary Figure 3B**). Gaussian noise did not elicit obvious plateauing, which is as expected as its independent additive nature makes its effects in later stages much less marginal (**Supplementary Figure 3C**).

Gaussian blur, however, provided the clearest practically relevant example of sensitivity-limited premature plateauing (**Figure 3D**). Most metrics changed strongly during approximately the first 10 iterations, when the highest-frequency texture was removed, but subsequently flattened despite continued loss of multiple larger-scale intracellular blob-like structure. In several extreme cases of sensitivity-limited premature plateauing, such as with SSIM and foreground SSIM, metric values stabilized between 80% and 85% of their normalized fidelity range amidst ongoing intracellular texture decay. Preservation of object shapes seemed sufficient for earning the bulk of metric rewards, and destruction of subcellular texture incur only marginal penalty. Because excessive smoothing is a common failure mode of image predicting models, this metric behavior is particularly consequential.

For grid distortion and dilation, most metrics exhibited a pronounced initial response during the first 10–20 iterations, corresponding to the early introduction of geometric warping that visibly disrupted cellular alignment and local continuity (**Figure 3E; Supplementary Figure 3D**). Beyond this initial phase, the metrics diverged into two broad regimes. PSNR and foreground PSNR tended to plateau relatively early, with only marginal additional change despite continued increases in distortion strength. In contrast, DISTS, LPIPS, SSIM, and foreground SSIM continued to respond over a longer portion of the trajectory, tracking progressive deformation of cellular shapes and loss of spatial coherence. However, these metrics with prolonged responses still retain misleadingly high normalized scores of around 0.6 despite cells having been rendered as textureless squares. The foreground SSIM stood out as a partial exception to this pattern. By restricting evaluation to biologically relevant regions, it combined sustained response and highest empirical dynamic range of close to 0.80 against grid distortion.

Taken together, our analysis of the metric absolute response exposes a largely invisible limitation of conventional metric reporting. Most evaluated metrics will correctly place near-identical images close to their theoretical optimum, yet compress the subsequent transition from intact to severely corrupted images into only a fraction of their nominal range.

### Characterizing IQA metric behavior against real virtual staining predictions

Our simulated degradation experiments showed that commonly used IQA metrics have limited sensitivity to image degradation and are strongly influenced by variations in biology. We therefore next asked whether these limitations remain evident in realistic virtual staining settings with heterogeneous reconstruction errors, and unknown severity likely driven by biological domain shift and other technical factors. Our main goal here is to empirically characterize what metrics can and cannot reliably reveal about virtual staining generalization to OOD data.

To enable this analysis, we constructed PedCP90VS, the virtual staining derivative of PedCP90. Using these data, we trained three convolutional neural network (CNN)-based virtual staining architectures: vanilla U-Net as a baseline (3,4); WGAN-GP (2); and finally, a simplified 2D UNeXt architecture (13). These architectures represent the evolution of image-to-image translation in cell microscopy virtual staining. We restricted model training to U2-OS images from PedCP90VS plate 1, predicting from brightfield images each of the five Cell Painting fluorescence channels. Models attain stable convergence under the combined MAE and multi-scale SSIM objective within the 300 epochs, with more complex models converging at later epoch numbers (**Supplementary Figure 4A-B**). See **Methods** for more details on model training and optimization. We then generated predictions for the remaining PedCP90VS evaluation images, yielding 18,818 crops spanning held-out U2-OS and unseen cell-line contexts (**Supplementary Table 4**). Unlike PedCP90Sim, where degradation type and severity are explicitly controlled, PedCP90VS tests metric behavior when reconstruction errors arise naturally from learned models and neither their magnitude nor failure mode is known.

#### Biological content explains over 50x more metric variation than model architecture

We evaluated PedCP90VS predictions against their corresponding Cell Painting references using the same metrics as in the simulated-degradation analysis (**Figure 1C**). To determine what drives variation in metric scores, we performed ANOVA (type-II) incorporating target channel, evaluation cell line, evaluation seeding density, model architecture, training seeding density, and their interactions. We repeated the complete inference and analysis workflow as PedCP90Sim across four evaluation subsets spanning progressively stronger extent of OOD: held-out U2-OS, evaluation U2-OS from a separate plate, unseen cell lines from plate 1, and unseen cell lines from plate 2. Given reported successes with virtual staining in label-free microscopy, if the metrics are truly sensitive to reconstruction quality, terms involving model architecture and training conditions should explain appreciable metric variation. Underwhelmingly, model architecture and training seeding density together explained less than 3% of the variance in held-out U2-OS predictions and less than 1% of the variance across unseen evaluation cell lines (**Figure 4A-C**).

**Figure 4:**
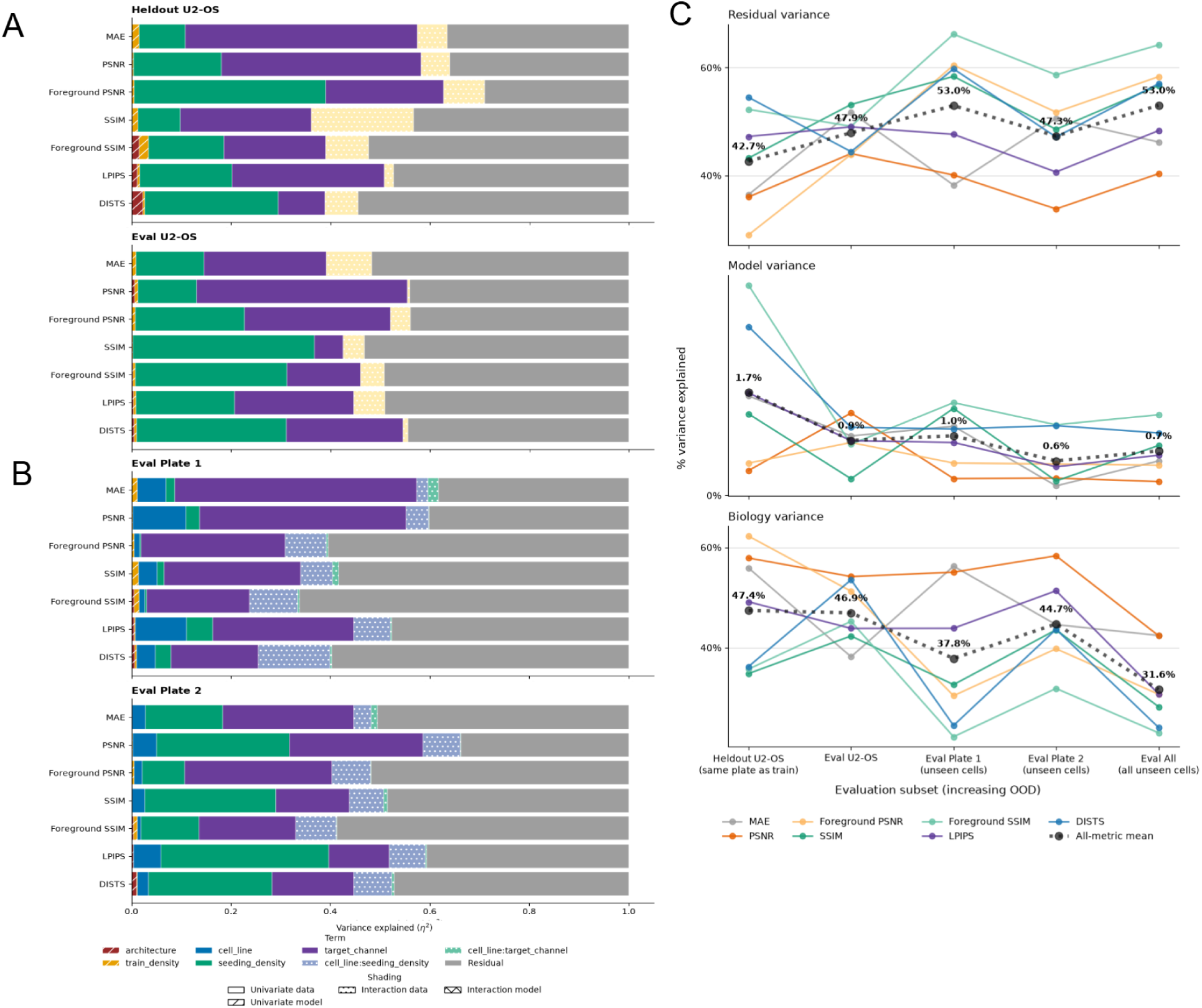
Metric response in realistic virtual staining evaluation setting. (**A**) Variance decomposition analysis attributing variation in heldout (top) and evaluation (bottom) U2-OS. (**B**) Variance decomposition analysis attributing variation in evaluation metric values in plate 1 (top) and plate 2 (bottom) unseen cells (**C**) Line plot summarizing the change in relative proportion of contribution by residual, modelling choice and biology.

In contrast, biological and imaging context accounted for a much larger fraction of metric variation. Terms associated with cell identity, seeding density, and target channel collectively explained approximately 30–60% of variation in held-out U2-OS, 35–50% in evaluation U2-OS, 20–55% in unseen plate-1 cells, and 30–60% in unseen plate-2 cells (**Figure 4A-C**). Mean normalized DISTS, LPIPS, SSIM, and foreground-SSIM scores testifies that the contribution of confounding effects revealed a cell-line specific systematic effect. Cell lines sourcing from osteosarcoma cancer or sharing mesodermal origin with U2-OS, such as G292, G401, and G402 received higher mean scores. Cell lines of neuroblastoma or of epithelial origin, such as SK-N-AS, KP-N-YN, SH-SY5Y, and NB-1 tended to score lowest (**Supplementary Table 4**). The cell line dependent mean metric score suggests that an appreciable portion of confounding might source from systematically easier or harder to predict cells.

An additional limitation became increasingly apparent with the stronger extent of OOD. Residual variance accounted for approximately 20–55% of score variation even in held-out U2-OS and reached 40–70% in unseen plate 1 cells, generally increasing as evaluation data moved farther from the training distribution (**Figure 4C**). Thus, under realistic virtual staining errors, a substantial and growing fraction of metric variation could not be attributed to either the modeled biological variables or the model-design factors under comparison.

Together, these results recapitulate the central limitation identified through the controlled degradation analysis: IQA scores are strongly shaped by biological context while providing little variation attributable to model architecture or training condition. As evaluation moves farther from training data, increased residual variation presence further weakens the interpretability of absolute metric differences. Consequently, these metrics may only reveal broad cell line-dependent trends, providing limited support for ranking architectures or attributing differences to model generalization.

#### Not all cell lines are equally generalizable by virtual staining models trained using U2-OS

The increasing residual variance observed under stronger extent of ODD suggested that reconstruction quality and systematic influence of biology were both becoming less measurable by the metrics. We therefore qualitatively inspected predictions across evaluation contexts to determine if and what forms of reconstruction heterogeneity accompanied this unexplained variation. All three architectures learned recognizable structure when evaluated near the training distribution in U2-OS cells. Visual quality of prediction became less consistent within and between the same architectures, however, as evaluation moved to unseen cell lines and cell density regimes (**Supplementary Figure 5**). Across architectures, some out-of-distribution images showed loss of texture, altered morphology, or incomplete recovery of fluorescence structure despite plausible overall appearance. The severity and form of these failures varied across biological contexts, suggesting models trained on U2-OS generalize unevenly across pediatric cell lines, with some biological contexts remaining comparatively stable and others producing heterogeneous, image-specific failures. This increasing within-context heterogeneity provides a qualitative counterpart to the growing residual variance observed in the metric analysis and highlights an additional challenge for summarizing out-of-distribution performance with aggregate scores alone.

#### Similar metric scores can correspond to qualitatively different modes of reconstruction failure

We next asked whether the variation captured by IQA scores faithfully represented these heterogeneous reconstruction errors. We first inspected representative Mito channel predictions from G292, G401, G402, U2-OS, and Saos-2 at a seeding density of 4,000 cells, selecting the three highest-, median-, and lowest-scoring predictions within each context according to normalized, direction-aligned DISTS (**Supplementary Figure 6A-E**). These cell lines occupied the favorable end of the DISTS distribution, with mean normalized scores of approximately 0.80–0.85 and relatively narrow empirical ranges of approximately 0.15–0.23 (**Supplementary Table 4**). Within individual cell-line contexts, DISTS generally preserved a useful ordering: higher-scoring predictions more often recovered the overall cellular silhouette and prominent mitochondrial signal, whereas lower-scoring predictions showed increasing intensity mismatch, incomplete structures, or spatial misalignment.

We next examined SK-N-AS, KP-N-YN, KNS-42, NB-1, and SH-SY5Y, which occupied the less favorable end of the mean metric score, with mean normalized DISTS values of approximately 0.59–0.65 (**Supplementary Table 4; Supplementary Figure 7A-E**). These predictions exhibited substantially greater qualitative heterogeneity. Failure modes included hallucinated fluorescence, checkerboard-like artifacts, excessive signal amplification, widespread attenuation, loss of puncta, and incorrect subcellular localization. DISTS broadly separated relatively plausible predictions from severely corrupted examples, but the same score interval could contain fundamentally different types of failure. Predictions near 0.4–0.5, for example, could reflect either invented fluorescence, severe illumination mismatch, or structured artifacts (such as loss/gain/misalignment of subcellular structure), where predictions near 0.8 could still contain conspicuous errors in mitochondrial localization or texture.

Taken together, our findings in real virtual staining predictions identify consistent metric limitations as simulation: IQA scores reflect coarse reconstruction fidelity, but they cannot by themselves establish equivalent biological fidelity or fully characterize out-of-distribution failure. Reliable virtual staining training and evaluation with IQA metrics therefore requires collecting and examining score distributions between and across wide variations in biological contexts, complemented by manual inspection of best case, average case and worse case predictions.

## Discussion

Reference-based IQA metrics are deeply embedded in virtual staining evaluation for microscopy, and are largely assumed as stable, unbiased numerical measures of reconstruction fidelity (16,23). They are also currently the best options available for evaluation. In a comprehensive analysis evaluating the stability and bias of these metrics, we find that these metrics should be doubted. In controlled degradation experiments, we found that metrics reflected the biological content of the image being evaluated more so than the actual virtual staining performance. Their absolute numerical scales were poorly utilized, with severe degradation compressed into a small fraction of the nominal score range and several metrics becoming weakly responsive while recognizable image information continued to disappear. When applied to real virtual staining predictions, biological context again explained substantially more metric variation than model architecture or training condition, and similar metric values corresponded to qualitatively different reconstruction failures. The central limitation of current IQA-based evaluation is therefore not simply insufficient sensitivity, but the absence of a one-on-one mapping between metric value and biological reconstruction fidelity.

Some metrics behaved more favorably than others. DISTS, for example, provided the strongest overall compromise between degradation sensitivity and resistance to biological confounding among the metrics evaluated, particularly for difficult spatial perturbations. Foreground-restricted SSIM and PSNR by the Spotlight method, offered a complementary advantage by reducing dependence on foreground occupancy, seeding density, and cell-line identity. These findings support deliberate metric selection rather than interchangeable use of conventional IQA scores. We do not, however, identify them as universally-reliable metrics as they remain dependent on biological context and degradation type.

The combination of biological confounding, restricted empirical range, and failure-mode dependence is particularly consequential for model comparison. A metric may assign different scores to comparable reconstruction errors because the underlying cell type, channel, or confluence differs from training; conversely, substantially different reconstruction errors may occupy nearly the same region of its numerical scale. The metric response can further change with the type of reconstruction failure modes being measured, such that the same metric would differentially penalize blurring and inconsistent shapes. Consequently, neither a favorable absolute score nor a small difference between competing models has a universal interpretation. Therefore, genuine model differences may be small, poorly captured by the selected metrics, obscured by biological variation, or expressed through failure modes that the metrics weight differently. Either way, progress driven by ever more sophisticated architectures is constrained either by flawed evaluation metrics, or by the marginal size of architecture-driven gains.

These limitations narrow the claims that IQA metrics can support. Current metrics remain useful for detecting particularly unstable evaluation conditions, and summarizing reconstruction similarity within carefully matched biological contexts. Their interpretation is safer when model predictions are in a closely related biological context. Even these practices, however, mitigate rather than resolve the underlying problem. An ideal virtual-staining metric would respond strongly to biologically-meaningful reconstruction errors while remaining invariant to irrelevant variation in cell type, cell number, fluorescence abundance, morphology, acquisition condition, and other properties of the reference image. The structures that distinguish biological contexts are likely inseparably convoluted with absolute measure of reconstruction quality. A metric cannot simply become insensitive to these features without risking insensitivity to errors.

The challenge identified here is not unique to virtual staining of cell microscopy or histopathology. Other imaging fields have repeatedly confronted the mismatch between generic similarity scores and the quantities that ultimately matter. In natural-image restoration, perceptual-similarity benchmarks have explicitly compared candidate metrics against human judgments, motivating learned measures such as LPIPS and evaluation frameworks such as the PIRM challenge that jointly considered distortion and perceived image quality (24,25). Collectively, these efforts establish an important precedent: evaluation metrics should themselves be validated against the observer, decision, or scientific task for which the reconstructed image will be used. Our findings extend this principle to fluorescence virtual staining by showing that, even before downstream utility is considered, commonly used IQA metrics can be systematically modulated by biological context and can respond inconsistently to distinct reconstruction failures.

For this reason, development of increasingly sophisticated scalar IQA metrics may not be the only, or ultimately the most informative, path forward. Virtual staining is generally developed to enable a biological measurement or discovery, not to reproduce fluorescence images for their own sake. Evaluation can therefore be moved closer to the intended application. In a drug screen or perturbation experiment with paired fluorescence ground truth, for example, virtual stains can be evaluated by asking whether they preserve known treatment-associated phenotypes, perturbational relationships, feature profiles, dose responses, or separability between biological conditions. PaPIS, for example, introduces an application aware, perceptual similarity metric for virtual histopathology moving away from reconstruction fidelity alone (26). Tonks et al. introduced information gain as a cell-wise, strictly proper scoring rule for cell microscopy virtual staining feature preservation (20). Such assessments provide a reference for the biological variability that the model is expected to recover and directly test whether conclusions drawn from virtual stains agree with those drawn from experimentally measured fluorescence.

We therefore advocate a shift from metric-centric validation toward biologically-anchored evaluation of virtual staining models. IQA metrics are still necessary and needed as supervision objectives and surrogate metrics for model hyperparameter optimizations. But claims of model generalizability and performance should ultimately be supported by evidence that the virtual stains preserve the biological signals for which they are intended. Such a framework should not require a context invariant IQA metric. Instead, it should ground model validity by asking direct application-relevant questions: whether replacing experimental fluorescence with a virtual stain preserves the scientific conclusions of the image based profiling experiment, for example.

### Study limitations

This study has several limitations that define the scope of its conclusions. The controlled-degradation suite covered a limited set of manually specified transformations. Although these transformations represent distinct intensity, texture, and noise, they do not reproduce the full complexity of virtual staining errors. We did not combine degradations, for example. The morphology operations are limited to even, constant effects that erode or dilate grayscale objects as a whole. Realistic virtual staining artifacts most likely contain mixtures of spatially heterogeneous textural and morphological errors specific to the channel and biology that cannot be represented by a single degradation parameter or even the full suite of degradation combined. We also selected transformation severities to produce interpretable trajectories rather than calibrated to the prevalence or biological consequences of errors produced by deployed models.

Furthermore, to isolate metric trajectories from biological variation, we intentionally restricted the empirical dynamic-range and saturation analysis to one field of view from a single cell line, fluorescence channel, and seeding-density condition. The resulting curves provide illustrative evidence of scale compression but should not be interpreted as universal calibration functions. Empirical ranges and saturation points may differ across channels, cell types, image intensity distributions, preprocessing procedures, and acquisition systems. Additionally, we did not directly quantify the relationship between the synthetic transformations and architecture-specific prediction failures (e.g. does the former well represent the latter; is the latter merely a mixture of former). The controlled experiments establish how metrics respond to known perturbations, but they do not determine which perturbations best approximate the residual errors of U-Net, WGAN-GP, or UNeXt predictions. Future work could decompose real prediction residuals into empirically derived error classes or construct perturbations from observed model failures. This would permit more direct evaluation of whether a metric is sensitive to the errors that actually distinguish competing architectures.

The biological and technical diversity of our dataset was limited to one single Cell Painting dataset collected over a single small batch, on a relatively limited selection of pediatric cancer cell lines, and free of any perturbations. This design allows controlled comparison across substantial biological heterogeneity but will not simulate data distribution drifts across laboratories, microscopes, staining protocols, treatment perturbations, or acquisition platforms. Our virtual staining models were also trained from single-plane brightfield images, which may be an overly limited and non-robust source of label-free information for prediction of Cell Painting fluorescence, as discussed by cell microscopy virtual staining work (2,13). Incorporating additional information such as brightfield z-stacks, phase-reconstructed inputs, or deconvolved volumetric images could reduce this information bottleneck and provide the model with more complete morphological cues from which to infer fluorescence structure. Furthermore, nominal seeding density imperfectly represented final confluence because proliferation, attachment, and survival differed among cell lines. Residual confounding by actual cell count and foreground occupancy is therefore likely.

Our evaluation was also restricted to seven of the most commonly-used full-reference IQA metrics and two foreground-masked variants. We did not assess all available perceptual, population statistic based, and segmentation based methods of image quality assessment (**Supplementary table 2**) (27). We likewise did not perform formal expert perceptual scoring or comprehensive feature-based evaluation of Cell Painting profiles (28). The present results therefore do not establish that all possible evaluation methods share the same limitations. Instead, they show that several widely used scalar reference metrics do not provide a sufficiently context-invariant evaluation scale on their own.

Finally, the virtual staining model evaluation included three convolutional architectures trained with a common objective and optimization configuration. It did not include transformer-based, diffusion-based, or other emerging generative architectures. Each configuration was represented by a single training run without extensive architecture-specific hyperparameter optimization or repeated random seeds. Consequently, the study was not designed to estimate the maximal performance of each architecture, characterize training uncertainty, or establish definitive architectural rankings. These limitations reinforce, rather than weaken, the decision not to interpret small metric differences as proof of model superiority.

## Methods

### PedCP90 Dataset processing

We analyzed a pediatric cancer dataset of 18 cell lines, five cell seeding densities, collected at a 24 hour incubation period across two plates. We processed these data with a CellProfiler pipeline to apply illumination correction, single-cell segmentation, and feature extraction to generate two plate-specific SQLite files (29). For details about the CellProfiler pipeline, see https://github.com/WayScience/pediatric_cancer_atlas_profiling. We then used CytoTable to process the SQLite files into parquet files and CoSMicQC to perform single-cell segmentation quality control (30,31). We extracted QC’d center x, y coordinates to identify where to crop single cells. The full PedCP90 dataset post-QC contains 10,248 full FOV images from 380 unique wells imaged.

### PedCP90Sim creation

The PedCP90Sim dataset is obtained by applying the degradation suite on a subsetted and cropped derivative of the raw PedCP90 dataset, which contains full FOVs (1080 by 1080 pixel images).

We first describe the cropping process: Cropping is configured to be 256 by 256 windows, and their centers are directly informed by the center coordinates of the CellProfilier nuclei segmentation of the full PedCP90 dataset. This initial cropping step resulted in 46,289 nuclei centered crops. As the quality control step on the original PedCP90 removes full FOVs falling below a certain quality threshold, there is an uneven distribution at both the full FOV and crop level across biological conditions. To ensure balanced representation by all 90 conditions at the crop resolution, we randomly sample 50 crops from PedCP90 to max 50 full FOVs per well. The end result is a cropped dataset containing 18,818 unique crops each of dimension 256 by 256 and with five Cell Painting channels. All crops and all channels are uniformly normalized by the image bit depth (16) so all pixel values post normalization are between 0-1.

With the crop dataset obtained, we apply the following suite of image degrading transforms to all five channels of every crop. The transform functions are either directly from the package *albumentations* (*GaussNoise, GaussianBlur, RandomGamma, GridDistortion*) or are custom implementations using *opencv* (*erode, dilate*) (32,33). Specific configurations are as **Supplementary table 3.**

The total number of degraded variants is the sum of parameters in parameter sweep across all degradation types. In this manuscript, we have 6 transform types x 6 parameters each = 36 degraded variants per channel and crop. Because most albumentations transforms have internal random states and are non-deterministic, to ensure reproducibility for this analysis, we re-initialize a new instance of these transform functions per crop and channel, all seeded by a master random seed of 42 which is further salted by the hash of the string values of the biological condition variables (cell line, seeding density, channel) and crop index. The end result of applying the degradation suite across the crop dataset is PedCP90Sim, containing 18,818 x 36 degraded crops each with 5 channels. Every degraded crop can be mapped back to one and exact one ground truth fluorescence crop.

### Simulated degradation analysis with PedCP90Sim

#### PedCP90Sim metric evaluation

We evaluated four full reference metrics: SSIM, Foreground SSIM, PSNR, Foreground PSNR, DISTS, LPIPS, MAE on the PedCP90Sim dataset. Specifically, one metric value is obtained per unique combination of metric, channel, and degraded crop, against the matching ground truth crop. This resulted in 7 metric x 5 channels x 18,818 crop x 36 degraded variant metric values for a total of 23,710,680 values. Each value is associated with 1) cell line 2) seeding density 3) channel 4) metric 5) origin plate and well metadata. We used this dataset in downstream analyses.

#### PedCP90Sim ANOVA analysis

Variance partitioning is achieved through ANOVA (type II). Specifically, we fit one type II ANOVA model per combination of metric and degradation, which includes all variability in biology, channel and severity. We model the metric values against four univariate terms: severity, cell line, seeding density, and channel; three interaction terms between biology: cell line:seeding density, cell line:channel, and seeding density:channel; three interaction terms between degradation severity and biology: severity:cell line, severity:seeding density, and severity:channel. We compute the variance explained percentage associated with each univariate or interaction term as the fraction of term sum of squares over total sum of squares. The residual component is computed as the complement of all summed explained percentages.

This full analysis fitted 6 degradation type x 7 metric = 42 ANOVA models, each having an input sample size into each ANOVA fit is therefore 5 channels x 18, 818 crop x 42 metric-degradation pairs against 10 terms.

#### PedCP90Sim nested regression analysis

To more directly compare metric sensitivity to controlled image degradation against metric dependence on biological context, we performed a nested regression analysis complementary to the global ANOVA variance partitioning. Analyses were stratified by fluorescence channel to remove channel-associated differences in metric distributions and were performed separately for each combination of degradation type and IQA metric.

Within each combination, we evaluated cell-line identity and seeding density as biological covariates in separate analyses. For each analysis, we first fit a restricted linear model

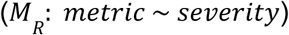

in which metric value was modeled as a univariate function of degradation severity alone with an intercept term. We then fit two corresponding full models

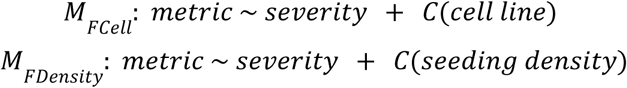

by adding either cell-line identity or seeding density as a categorical predictor. The coefficient of determination of the restricted model, referred to as restricted R^2^ in this manuscript, was used to quantify the degree to which metric variation was explained by the known magnitude of image degradation. The additional explanatory contribution of biological context was quantified using partial R^2^, computed from the full model via

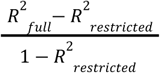

Thus, restricted R^2^ represents metric sensitivity to the intended degradation signal, whereas partial R^2^ measures the additional influence by seeding density or cell line, after accounting for degradation severity.

To estimate the stability of these R^2^ quantities, each nested-model comparison was repeated across 300 bootstrap resamples generated by sampling observations with replacement at the original sample size. Biological groups containing fewer than 25 observations were excluded from fitting, and bootstrap sampling used a fixed random seed of 42 for reproducibility. For each metric, degradation type, fluorescence channel, and biological covariate, we summarized restricted and partial R^2^ by their bootstrap mean and 95% percentile interval. Across seven metrics, seven degradation types, five fluorescence channels, and two biological covariates, this produced 490 nested-model comparisons. To provide a compact descriptive comparison of biological-context dependence relative to degradation sensitivity, we additionally calculated a metric burden statistic as the ratio of partial R^2^ to restricted R^2^, with smaller values indicating metrics whose responses were more strongly associated with controlled degradation severity relative to biological context.

#### Ceiling-attainment and empirical dynamic range analysis

To characterize the absolute numerical response of each IQA metric to progressively increasing image degradation, we first placed metric values on a common normalized fidelity scale in which 1 represents the specified upper bound corresponding to highest image fidelity and 0 represents the lower bound. Defining this scale required specifying a reference numerical range for each metric. MAE is naturally bounded between 0 and 1 for images normalized to this intensity range. SSIM is mathematically bounded between -1 and 1; however, strongly negative SSIM values indicate substantial structural anticorrelation and were not encountered as a meaningful regime in the present image comparisons. We therefore clipped SSIM values to [0,1] and used this interval as its reference range. PSNR is non-negative but has no finite theoretical upper bound because it approaches infinity as reconstruction error approaches zero. We therefore used 80 decibels (dB) as an operational upper bound for PSNR normalization. This value provides a finite reference ceiling while remaining substantially higher than PSNR values encountered for degraded images in our analysis; normalized PSNR should therefore be interpreted relative to this imposed operational range rather than as a fraction of a theoretical maximum. Foreground-restricted variants used the same numerical bounds as their corresponding full-image metrics. Metrics whose native orientation is lower-is-better were inverted after normalization so that all analyzed trajectories shared the same interpretation: larger normalized values indicate greater fidelity to the reference image.

For each reference crop, we compared each progressively degraded variant with its corresponding undegraded image, producing an ordered metric trajectory from the lowest to the highest degradation severity. We summarized the absolute response of each metric to each degradation type using two complementary properties. Ceiling attainment quantifies the normalized fidelity score at the lowest non-zero degradation severity and therefore measures how closely a minimally perturbed image approaches the nominal maximum score. Values approaching 1 indicate that small perturbations receive scores close to perfect fidelity, whereas lower values indicate substantial numerical penalties even for minimally degraded images. Empirical dynamic range quantifies the fraction of the specified metric range traversed over the complete degradation trajectory and was calculated as the difference between the maximum and minimum normalized fidelity scores observed across degradation severities. Because all normalized metrics span a reference range of [0,1], empirical dynamic range also directly represents the proportion of the available numerical range occupied by the experimentally observed degradation response. A larger empirical dynamic range indicates greater numerical separation between mild and severe degradation, whereas a restricted range indicates that substantially different image-quality states are compressed into a comparatively narrow interval of metric values.

### PedCP90VS creation

#### Train data preparation

We assigned all U2-OS images from the first plate to training, whereas images from the 17 unseen cell lines and an independently acquired U2-OS plate were reserved for evaluation. We further divided the U2-OS training images according to five seeding-density levels, producing five density-specific training datasets. Evaluation images were similarly stratified by cell line and seeding density, yielding 90 biological evaluation conditions comprising 17 out-of-distribution cell lines and a held-out U2-OS replicate across five seeding-density levels

Paired brightfield and Cell Painting fluorescence images were used to train and evaluate label-free virtual staining models. Each field of view contained a single focal-plane brightfield image paired with fluorescence images corresponding to the Cell Painting channels DNA, RNA, ER, mitochondria, and AGP. Images were annotated by cell line, assay plate, and seeding-density condition.

Because differences in confluence can strongly affect the foreground-to-background ratio of randomly sampled image crops, we used CellProfiler-derived segmentation masks to guide crop selection in both train and evaluation dataset. Specifically, non-overlapping 256 × 256 pixel crops were selected only when they contained at least one segmented nucleus, ensuring that training and evaluation crops were enriched for cellular foreground rather than dominated by empty background.

Additionally, because the same number of wells and FOVs were acquired for all seeding densities of the U2-OS train dataset, our segmentation based crop selection procedure naturally produced different sample sizes. All train datasets were augmented or downsampled to the uniform sample size of 10,000 crops to ensure all model training sees an equal number of image crops.

#### Model training

For each architecture, we trained separate models for each combination of training density and target Cell Painting channel, yielding a total of 75 virtual staining models. All models used the same optimization configuration: a learning rate of 0.0002 and a combined MAE and multi-scale SSIM objective adapted from Liu et al. (13). All models are trained for 300 epochs with a mini-batch size of 8, which helps us balance hardware demand and still give all models ample training iterations for convergence. Each trained model was then selected based on best validation performance over the 300 epochs, and evaluated on image crops of 1 by 256 by 256 containing at least one Cell Profiler segmented nuclei center, from all held-out biological conditions. The evaluation set consisted of 15 held-out cell lines across five seeding-density levels, yielding 75 prediction conditions per target channel. Predicted fluorescence images were compared with their matched ground-truth Cell Painting crops using the same set of seven IQA metrics characterized in our previous analysis: MAE, SSIM, PSNR, foreground SSIM, foreground PSNR, LPIPS, and DISTS.

All models were trained using the same optimizer configuration. Adam optimization was used with a learning rate of 1e-5, beta1=0.9, beta2=0.999, and no weight decay. All models were trained for 300 epochs alike and the best performing epoch weight based on the datasets were selected for the final evaluation.

The training loss is a combination of vanilla MAE loss and multi-scale structural similarity, which was reported by Liu et al to facilitate better convergence (13).

### Realistic virtual staining model evaluation with PedCP90VS

#### PedCP90VS ANOVA analysis

Variance partitioning is achieved through ANOVA (type II). Specifically, we fit one type II ANOVA model per combination of metric and degradation, which includes all variability in biology, channel and severity. We model the metric values against five univariate terms: cell line, seeding density, target channel, architecture, and train seeding density. We compute the variance explained percentage associated with each univariate or interaction term as the fraction of term sum of squares over total sum of squares. The residual component is computed as the complement of all summed explained percentages.

This full analysis fitted seven ANOVA models, one per metric each having an input sample size into each ANOVA fit is therefore 1,411,350l predicted crops.

## Data Availability

We present our code to reproduce this analysis in https://github.com/WayScience/virtual_staining_metric_analysis. We present code to train virtual staining models in https://github.com/WayScience/virtual_stain_flow. We present CellProfiler processing of the initial PedCP90 dataset in https://github.com/WayScience/pediatric_cancer_atlas_profiling.

## Acknowledgements

A portion of this work was funded by Alex’s Lemonade Stand Foundation ‘A’ Award and Tap Cancer Out (Grant #23–28306 to G.P.W.). We thank Jamie Cheah and Christy Biji for generating the PEDCP90 dataset, Erin Weisbart for facilitating its upload to the Cell Painting Gallery, and Jenna Tomkinson for CellProfiler processing. We also thank Cameron Mattson for contributions to the virtual staining training repository, and Michael Lippincott, Erik Serrano, Jenna Tomkinson, and Dave Bunten for code review.

## Supplementary Tables

**Supplementary Table 1.** *Cell Painting dataset*.

**Supplementary Table 2**. Metric selection and justification.

**Supplementary Table 3**. Image degradation suite parameters.

**Supplementary Table 4**. Mean metric score across cell line, seeding density, channel and architecture.

## Declaration of generative AI and AI-assisted technologies in the manuscript preparation process

During the preparation of this work, the authors used Claude Sonnet 5 for increasing clarity of certain sections of text and ChatGPT codex for select code contributions. The authors reviewed and edited the output as needed and take full responsibility for the content of the published article.

## Notes

https://github.com/WayScience/virtual_staining_metric_analysis

https://github.com/WayScience/virtual_stain_flow

https://github.com/WayScience/pediatric_cancer_atlas_profiling

### Competing Interest Statement

The authors have declared no competing interest.

