## Supplementary Figures for "On doubting image quality assessment metrics for microscopy virtual staining"

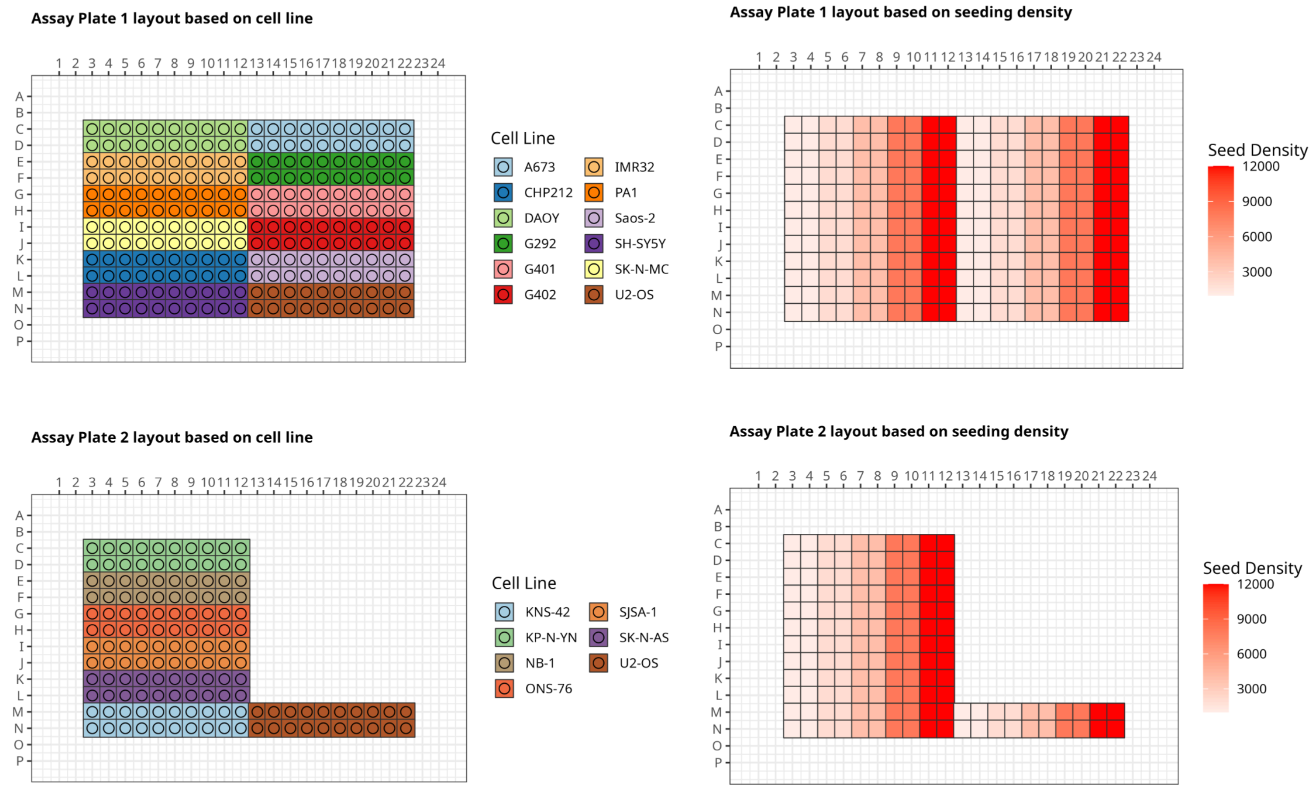

Supplemental Figure 1: PedCP90 Platemap.

Platemaps of the pediatric cancer imaging dataset. Eighteen cell lines were seeded at five densities (1,000, 2,000, 4,000, 8,000, and 12,000 cells/well), with four replicate wells per cell line–density combination, and imaged after 24 h of incubation across two 384-well plates. U2-OS was included on both plates using the same five seeding densities and replicate structure, providing a shared condition across plates.

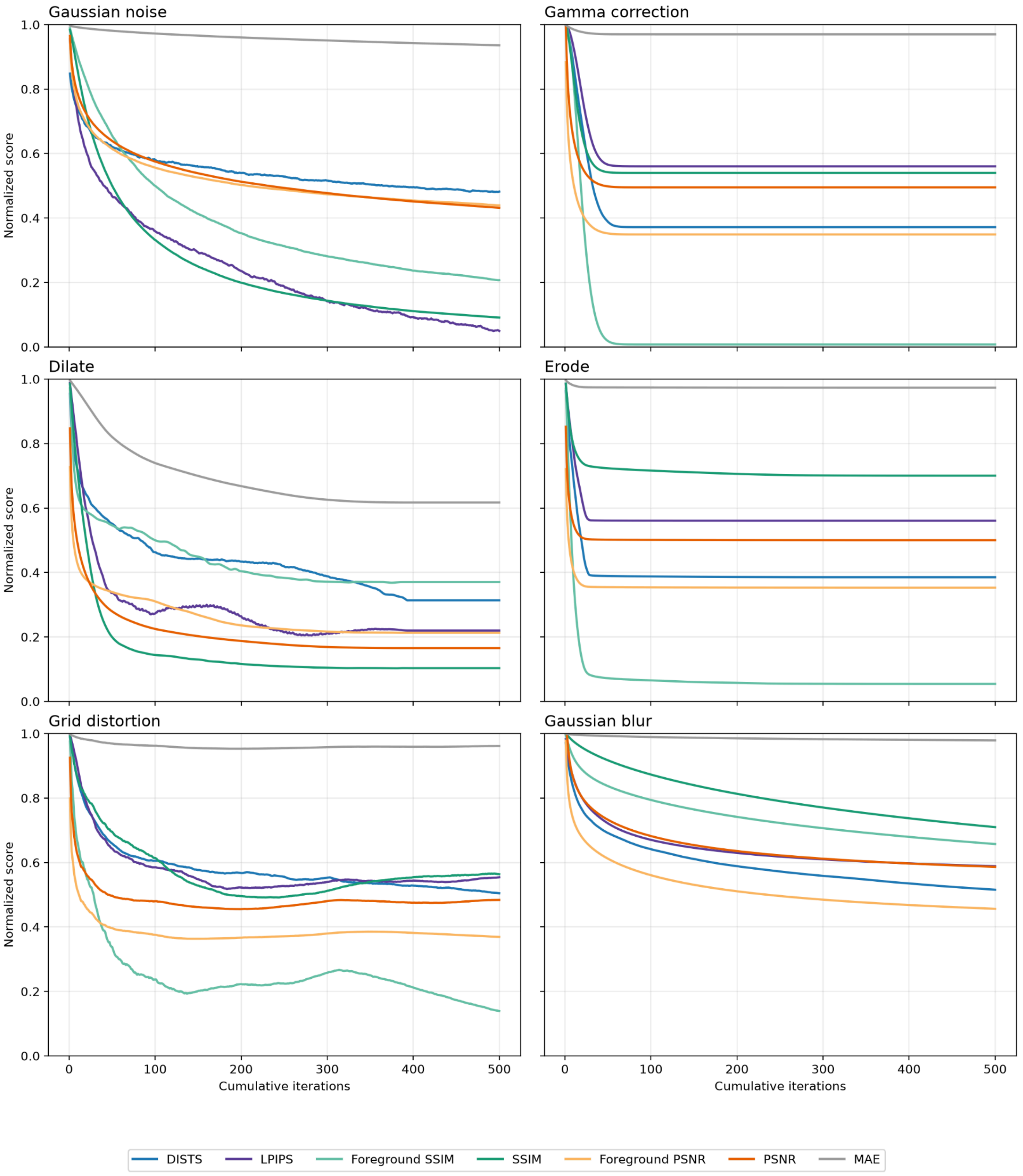

Supplemental Figure 2: Metric full absolute response curves.

Absolute responses from IQA metrics against 500 iterations applied image degrading transforms, metric values normalized by nominal range and direction aligned as higher is better. All metric response trajectories organized by degradation types for the full 500 iterations.

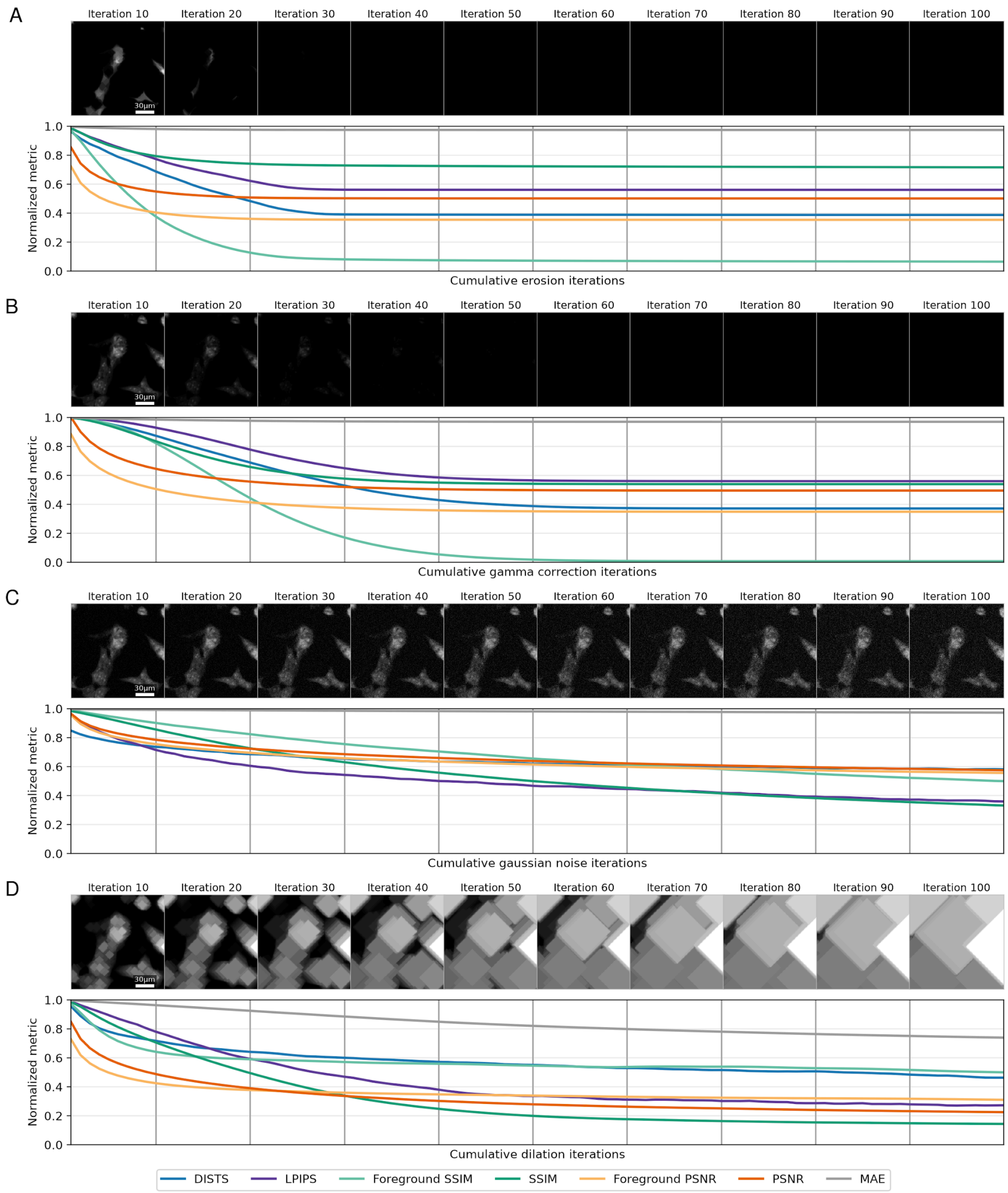

Supplemental Figure 3: Metric absolute response curve and montages, plateauing pattern corresponding to degradation in first 100 iterations.

In this supplemental figure we present the remainder of the metric plateauing responses corresponding to the content exhaustion pattern. In **A**) Incremental dimming gamma correction, and (**B**) Incremental erosion. In these examples, brighter pixels are multiplicatively made darker or assigned the minimal local neighborhood value. Both degradation modes cause the image to quickly become essentially blank, and therefore correspond to the reasonable, content-exhaustion induced plateauing response. (**C**) Incremental gaussian noise corresponds to a much less apparent metric plateauing response. This is expected as gaussian noise operates additively and independently from iteration to iteration, sprinkling the same amount of white noise each application, which results in sustained metric responses. (**D**) Incremental dilation elicits a similar sensitivity-limited premature plateauing response as the grid distortion example.

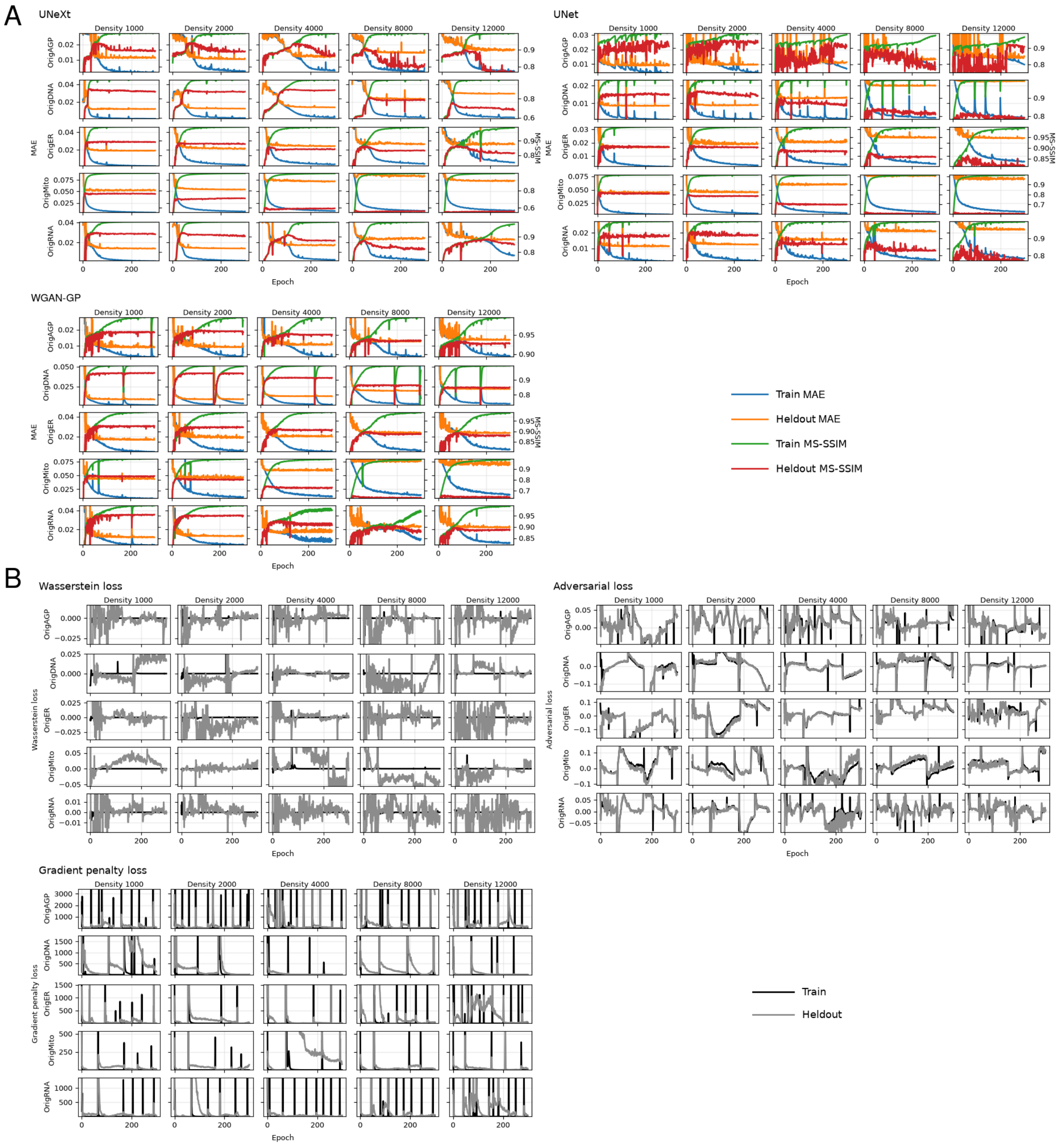

Supplemental Figure 4: Training and heldout (validation) loss curve across all 75 virtual staining models, full 300 epoch of training.

(**A**) Shared generator loss curves across the 3 architectures. The blue and orange lines correspond to train and heldout MAE loss, respectively. The green and red lines correspond to train and heldout multi-scale SSIM, respectively. All architecture and training conditions displayed convergence patterns within 300 epochs of training, with the exact time point of convergence depending on dataset, channel and model architecture. (**B**) Additional WGAN-GP training specific losses, train curve is colored black, heldout gray.

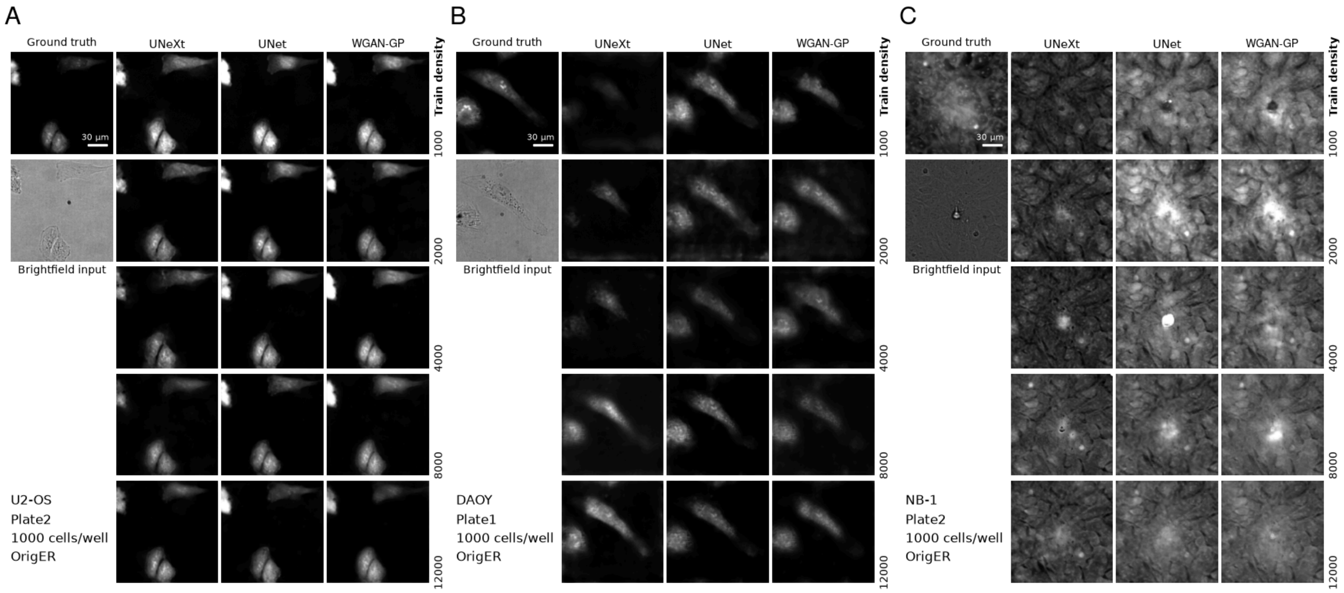

Supplemental Figure 5: Example predictions across differential OOD extent, all models, ER channel.

Predictions by all model architectures predicting the ER channel. Ordered by coarsely increasing extent of OOD. (**A**) Model predictions in eval U2-OS are quite fidel beyond altered brightness and remarkably uniform across all architectures and training seeding density. (**B**) Model predictions in DAOY, invention of dimmer objects and varied enlargement and shrinking of existing cells are apparent. (**C**) Model predictions in NB-1. The ground truth looks overly confluent already, probably due to fast proliferation of cells. Multi-cell tiling invention is common and extremely varied. Artifacting in the form of local brightening and dimming is also prevalent and varied.

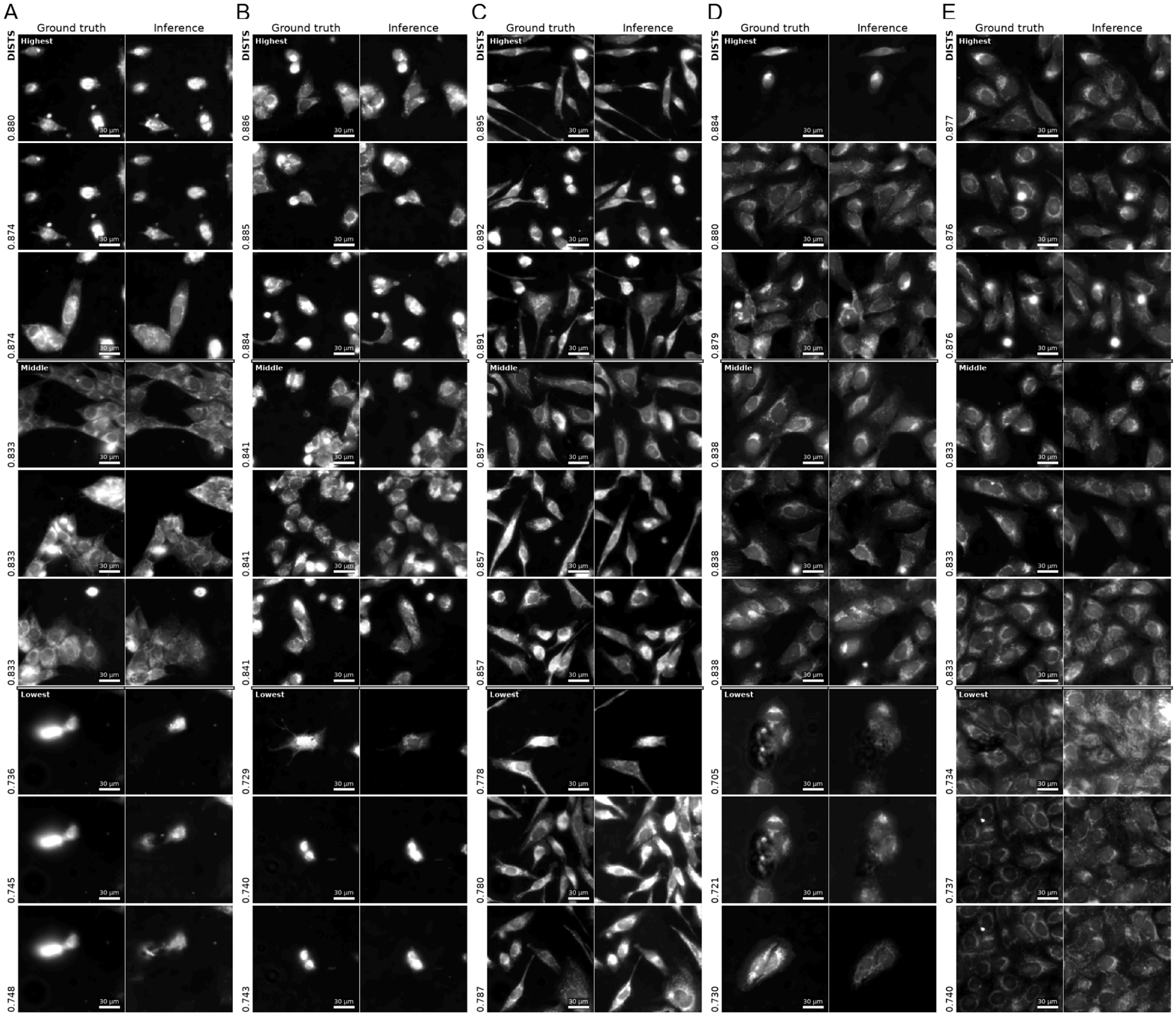

Supplemental Figure 6: Predictions receiving best, medium and worst DISTS metric scores per cell line across all models, less OOD cell lines. ER channel.

Representative good, mediocre and bad predictions according to DISTS, from cell lines more similar to train biology. All predictions presented next to ground truth. DISTS value received by each prediction is annotated at the left of each reference-prediction pair. (**A**) G292 (**B**) G401 (**C**) G402 (**D**) Plate 2 (eval) U2-OS (**E**) Saos-2.

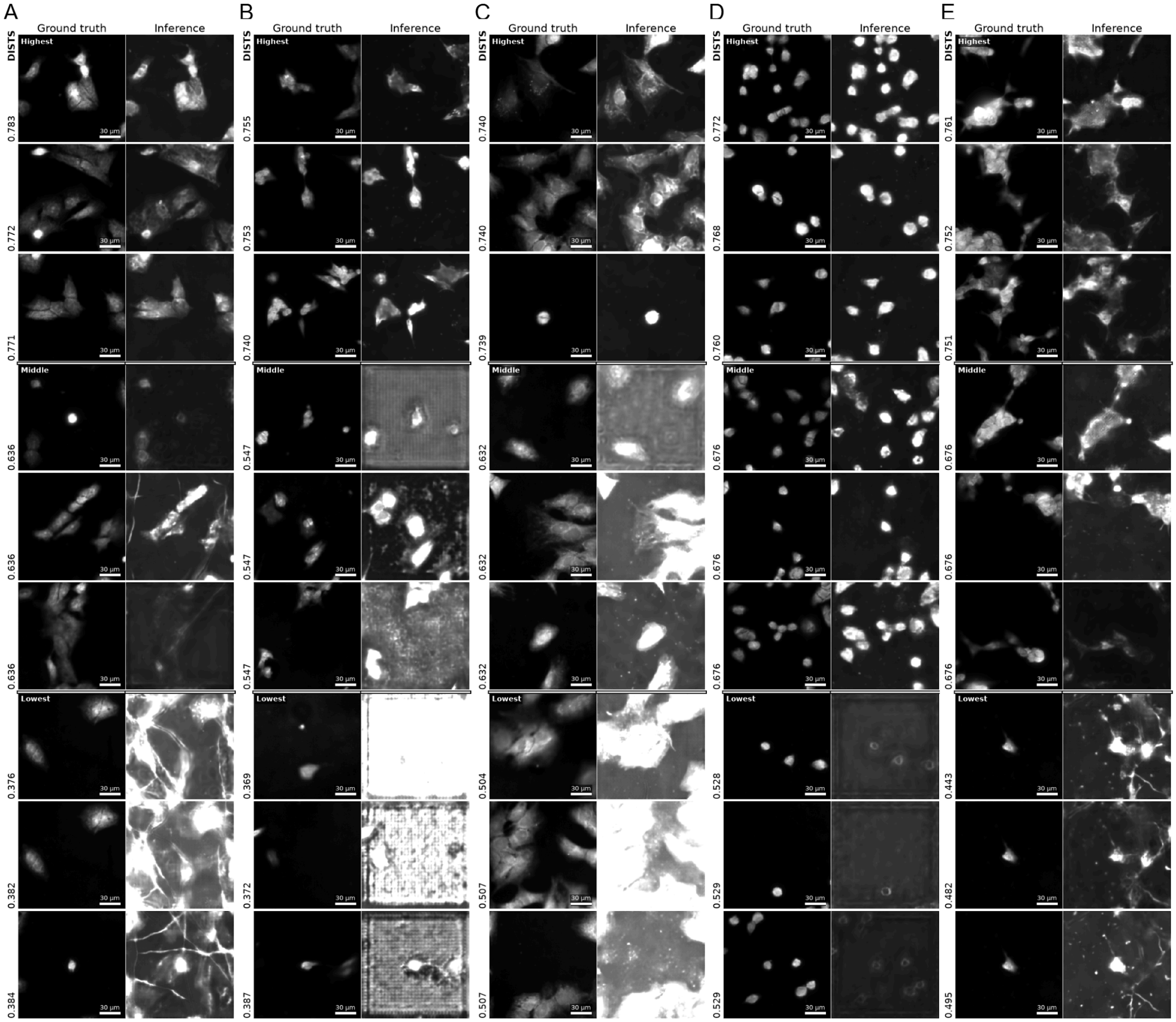

Supplemental Figure 7: Predictions receiving best, medium and worst DISTS metric scores per cell line across all models, more OOD cell lines.

Representative good, mediocre and bad predictions according to DISTS, from cell lines more dissimilar to train biology. All predictions presented next to ground truth. DISTS value received by each prediction is annotated at the left of each reference-prediction pair. (**A**) SK-N-AS (**B**) KP-N-YN (**C**) KNS-42 (**D**) NB-1 (**E**) SH-SY5Y.
